# Refining the resistance-to-damage (RD) score to predict operational Insecticide-Treated Net lifespan and identify paths to innovation

**DOI:** 10.1101/2025.03.02.641027

**Authors:** Frank Mechan, Stephen Poyer, Julie-Anne Akiko Tangena, Matt Worges, Eleanore Sternberg, Albert Kilian, Hannah Koenker, Amy Wheldrake, Fadi Junaid, Christen Fornadel, Stephen J. Russell, Angus Spiers

## Abstract

**Background:** Insecticide Treated Nets (ITNs) vary greatly in their physical characteristics yet the relationship with field durability remains poorly understood. To address this, we refined the existing resistance-to-damage (RD) metric for greater predictive power in estimating operational lifespan and to identify key avenues of innovation.

**Methods:** The original resistance-to-damage (oRD) score was calculated by comparing four physical attributes; snag strength, bursting strength, hole enlargement resistance and abrasion resistance; against aspirational values previously determined to provide resistance to routine damage. Here, the weighting of each is improved to provide a better prediction of field life, resulting in a weighted resistance-to-damage (wRD) score. Durability data from 37 field sites (across 14 countries) involving 10,000 campaign ITNs was used to quantify the relationship between physical characteristics and field life. Bivariable linear regression models were used to assess the correlation between RD scores and ITN survival time. A Cox regression survival analysis was applied to examine the impact of the RD score on physical survival, adjusting for net care and environment variables (Risk Index), evaluating the risk of ITN failure from the time they were first hung.

**Results:** The bivariate analysis, not adjusting for differences between sites, shows that the wRD was a better predictor for net lifespan than oRD (wRD r^2^= 0.07 oRD r^2^= 0.01). In the multivariate analysis, adjusting for risk index, the wRD model was able to explain more than 70% of variability in net survival (wRD r^2^ = 0.74). Consequently, when comparing nets with different wRD scores in the same site, a 10-point increase in wRD score was associated with a 3.9-month gain in median survival (*p*=0.016). Increasing wRD from 35 to 70, the range of current products, increases median field life by 13.65 months (95% CI: 3.03–24.16). An ITN product with an wRD score of 68 is associated with a median lifespan of 2.7 years. The increase in predictive power of the wRD score came primarily from increasing the weighting of the hole enlargement resistance variable, which is a strong differentiator between ITN products. In contrast, abrasion-resistance was a negligible contributor to predictive power.

**Discussion:** The wRD score more accurately predicts ITN survival in the field than the oRD score. These findings highlight that resistance to hole enlargement is the primary differentiating factor in predicting field life, with some products performing far better than others. Snag and bursting strength remain essential characteristics yet are similarly low compared to aspirational values across available ITN products. All ITNs WHO prequalified in 2020 perform poorly on snag strength, indicating this as a core bottleneck in achieving longer operational ITN lifespan. Consequently, resistance to snagging and hole enlargement should be keys targets for future innovation to enhance the physical durability of ITNs.

## Background

Insecticide Treated Nets (ITNs) are a core component of malaria vector control, presenting a physical and chemical barrier against the bites of *Anopheles* mosquitoes [1]. ITNs are proposed to be designed to be physically resistant against day-to-day use in the domestic environment, historically being expected to provide three years of operational lifespan [2]. National Malaria Programmes commonly distribute ITNs via mass distribution campaigns every three years, with timing set partly based on this expectation. However, recent modelling of ITN monitoring data across sub-Saharan Africa indicates that the median ITN retention is 1.64 years [3], reducing effective coverage until the next campaign (Commonly every three years). While ITN products vary in physical specifications, with different polymers being used across a range of fabric weights and knitting patterns [4, 5], there is no consensus or evidence base on which characteristics, if any, provide the longest operational use.

Owing to the high operational costs of malaria vector control programmes and heavily constrained funding environment, there has been a strong incentive to keep the price per unit of ITNs low [6]. This has made it difficult for more expensive but potentially more physically durable designs to enter the market, with even a reduction in ITN physical durability reported for the same products from 2013 to 2020[7]. There is now a groundswell amongst decision makers and ITN procurers to identify products that are the most physically robust in order to improve the coverage of ITNs in a cost-effective manner. Substantial improvements are required across the board if ITNs are to become more durable, which requires deeper understanding of the relationship between net physical characteristics and net lifespan.

While in principle ITNs could be made from a wide variety of available polymers and fabric constructions, the current crop of ITN products is either based on polyester (PET - polyethylene terephthalate) or polyethylene. Furthermore, all polyethylene nets use a tulle knitting pattern and all polyester nets a traverse pattern. Consequently, the currently available ITNs represent a very limited cross-section of what is technically possible. This may mean that a stepwise change in net design is needed to expand variability and physical net durability, which will be more expensive to produce. However a case could be made to fund more expensive designs if they were demonstrated to offer an extended operational life [8].

ITN products recommended by the World Health Organization Pesticide Evaluation Scheme (WHOPES) or, more recently, prequalified by the Vector Control Group of the WHO Prequalification Team (PQT-VC) were assessed in 2013 and 2020 respectively for their physical and design characteristics at the Nonwovens Innovation & Research Institute (NIRI) laboratories. Building on research aimed at understanding the sources of damage leading to holes in mosquito nets [9], the resistance-to-damage (RD) score was developed as a composite metric for quantifying the physical durability of an ITN product relative to an aspirational value [10-13]. The aspirational values were established based on analyses of human factors, considering the associated real-life forces applied to ITNs during normal use [7]. The original RD (oRD) score uses four performance indicators measured in the laboratory to evaluate an ITN’s resistance to routine stresses in the field; snag strength, bursting strength, hole enlargement resistance and abrasion resistance [10, 12, 13]. These performance indicators are included in the complete product dossier required for WHO prequalification assessment of ITNs and will be mandatory to report for all WHO-prequalified ITNs from 2025 [14]. A secondary analysis of ITN field durability by Killian et al (2021) demonstrated that the calculated oRD scores and survivorship in the field are correlated, such that ITNs with oRD scores >50 had an additional operational lifespan of 7 months (compared to those with oRD < 50) [15].

In this prior analysis for oRD, the four performance indicators were assumed to contribute equally to the overall RD score. However, recent analysis indicates some types of damage are more common than others, with abrasion contributing to just 3.7% of total hole area while enlargement of existing holes through laddering and unravelling being responsible for 18.1% of hole area [11]. With access to a larger field database than those available in 2021, when oRD scores were first compared with field data [10], and greater insight into hole formation, we investigated whether the RD metric could be refined to better predict operational lifespan by optimizing the weighting of its performance indicators based. We refined the oRD score using durability data from 13 ITN brands across 37 study sites in 15 sub-Saharan African countries to strengthen the evidence linking RD scores to the survival of ITNs in serviceable condition in the field.

## Methods

### Refining the Resistance-to-damage score

The net characteristics previously associated with resistance to routine damage in the field are summarised in Supplementary file 1 [7, 10-13, 15]. Four physical ITN fabric attributes, which have been previously linked to field life, were quantified to calculate the oRD score: snag strength, bursting strength, hole enlargement resistance and abrasion resistance (Table 1)[10]. The value of each performance indicator was assessed as a proportion of an aspirational value, representing the ideal score needed to endure day-to-day stresses in the field. The outcome of these four tests was combined in an algorithm to produce a single oRD score (out of a maximum of 100). Each of the four indicators are weighted equally, meaning they are implicitly assumed to be of equal importance for operational lifespan (Eq. 1).

**Table 1:** Description of performance indicators used to calculate RD scores.

| Indicator | Type of damage measured | Aspirational value |
| --- | --- | --- |
| Snag strength | Fabric catching on a solid protuberance | ≥200 N |
| Bursting strength | Fabric multidirectionally tensioned | ≥700 kPa |
| Hole enlargement resistance | An initial hole gradually tensioned | 100 (No Laddering, No Unravelling, No Tearing) |
| Abrasion resistance | Cumulative rubbing of fabric against a rough surface | 400 rubs |

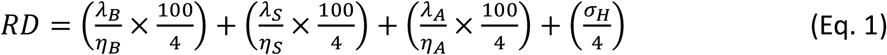

RD = Resistance-to-damage; λ_B_ = Actual bursting strength; η_B_ = Aspirational bursting strength; λ_S_ = Actual snag strength; η_S_ = Aspirational snag strength; λ_A_ = Actual abrasion resistance; η_A_ = Aspirational abrasion resistance; σ_H_ = Hole enlargement resistance score. Calculation can be done automatically using the RD score app [16].

### Correlating RD scores to field data

#### Data sources

ITN performance data came from durability monitoring studies conducted in 37 study sites across 15 countries in sub-Saharan Africa, representing 13 ITN brands. Primary data were collected by U.S. President’s Malaria Initiative (PMI)-funded studies conducted between 2015 and 2023, using standard protocols (Supplementary Table 2). We extracted site-level estimates of ITN physical survival as the median ITN survival time from the date of distribution. Estimates of median survival for each study site were determined using observations made at the 12-, 24- and 36-month timepoints. Confidence intervals around estimates were calculated using the Wald method. The median ITN survival time, from date of distribution, was 2.7 years. Given that some ITN use environments and individual behaviours could have a greater risk of damage to ITNs than others, we calculated a site-level risk index (RI) metric [14]. The risk index (RI) is a weighted average score based on variables collected during the baseline durability monitoring round. It reflects differences in environment factors, net handling behaviours, and individual net care attitudes across study sites (*manuscript in preparation*). As all inputs are percentages, the RI can vary from 0% to 100%, with higher values representing greater non-textile risks to ITN physical integrity. Variables included in the RI were selected based on hypothetical risk factors for net damage drawing on grey and published durability monitoring literature [14, 16]. ITN-level data were extracted from the same primary data source for survival analysis as described below.

The oRD scores were determined by NIRI and corresponded to the values presented by Wheldrake et al. [7]. The wRD score was calculated by the study team from the RD score components, obtained by NIRI. We used the 2013 published values for studies following ITN campaigns conducted between 2015 and 2018 and used the 2020 published values for study results from campaigns after 2018.

#### Analysis methods

We used bivariable linear regression models to assess the correlation between both the oRD score and wRD score compared to median ITN survival time at the site level estimated at the 12-, 24- and 36-month timepoints. We assessed model fit using the adjusted R-square value. We also performed a multivariate analysis adjusted for the presence of different environmental, household, and net-care variables by included the risk index in the model.

Having defined the wRD score, we assessed the association on wRD values and ITN physical survival using a Cox proportional hazards model with individual ITN data.

Data was set up for survival analysis as a duration format data set where each time interval for a net was a separate observation. Survival analysis was done using a per-protocol approach, i.e. risk of failure was considered to start only on the first observation where the net was found hanging. Failure was defined as a net reported to have been discarded by the household oris present but no longer in serviceable physical condition. Physical condition was defined according to the results of a hole assessment using the proportionate hole index (pHI), with serviceable nets defined as those with a pHI equal to or less than 642 (i.e. nets failed if the pHI was greater than 642). The time of failure was directly calculated from the reported time of loss by the respondent or taken as the mid-point between the last two surveys if time of loss was unknown. Results are reported as the gain in median survival in months for a 10-point change in the wRD.

Data were analysed in Stata v.18. Analysis of of the association between oRD, wRD and median ITN survial used study-site as the unit of analysis. Frequency weights were applied using the number of cohort nets remaining in the study at the 36-month data collection point as the weight (or 24-months for studies that ended after 24 months). Cox proportional hazards models used ITN as the unit of analysis, were unweighted, and included country as a random effect. For continuous variables, arithmetic means were used to describe the central tendency and the t-test for comparison of groups for normally distributed data. Otherwise, median and Kruskal–Wallis tests were used. Proportions were compared by contingency tables and the Chi squared test used to test for differences in proportions. For calculation of confidence intervals around estimates, the intra- and between-cluster correlation has been taken into account using the *svy* command in Stata.

Determinants of survival in serviceable condition after the net was first hung were explored using Cox proportionate hazard models. Factors were tested first in individual models which were then used to construct the final multivariate models. Final model fit was tested using a linktest and Schoenfeld residuals and log–log plots were used to check the proportionate hazard assumption.

## Results

### Refining the Resistance-to-damage score

Although in principle the oRD score is nominally equally weighted, in practice it is primarily made up of measures of resistance to initial hole formation (snag strength, bursting strength and abrasion resistance), with resistance to hole enlargement representing only 25% of the score (Eq. 1).

Assessing the performance of ITNs on the underlying measures highlights that all ITN products assessed perform poorly on snag strength relative to the aspirational value (Supplementary file 3A). While snag strength is indeed linearly related to fabric areal density (a proxy of the fabric weight), even the products with the highest areal density nets achieved only 25% of the aspiration. It was also observed that snag and bursting strength are highly correlated (supplementary file 4), yet high areal density is associated with good bursting strength performance compared to the aspirational value (supplementary file 5).

Resistance to hole enlargement contributes only 25% of the oRD score yet is directly responsible for the occurrence of large holes [7]. There is a notable difference in resistance to hole enlargement between tulle/monofilament polyethylene (PE) and traverse/multifilament Polyethylene terephthalate (PET), with PET nets consistently performing excellently on this metric yet PE nets varying greatly, and two examples of PET nets with low wRD (Supplementary file 3C). Mesh count was observed to correlate with performance on resistance to hole enlargement, with ITNs with the highest number of geometric structures per square inch performing best on this metric (Supplementary file 6).

While abrasion resistance was included as a performance indicator in the oRD when first developed, field data collected in years since from across sub-Saharan Africa shows that overall, only 3.7% of hole damage was caused by abrasion [11]. On this basis, we evaluate an alternative resistance-to-damage metric where abrasion is not considered. Furthermore, given that the same study observed most damage by hole area is caused by tearing of existing holes we doubled the weighting of hole enlargement resistance. Consequently, the wRD score was defined with the following weights: 25% snag resistance, 25% burst resistance, 50% hole enlargement resistance (Eq. 2).

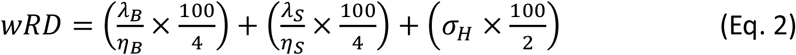

wRD = weighted resistance-to-damage; λ_B_ = Actual bursting strength; η_B_ = Aspirational bursting strength; λ_S_ = Actual snag strength; η_S_ = Aspirational snag strength; σ_H_ = Hole enlargement resistance score. Calculation can be done automatically using the RD score app [16].

#### Comparison of physical characteristics with oRD and wRD

The measured physical properties of all ITNs studied by Wheldrake et al [7] were collated to calculate both the oRD and wRD scores. Overall, we observed that wRD scores show a wider distribution than oRD scores, with nets clustering into two distinct groups: those with wRD scores of 30-40 and those with scores of 50-70 (Figure 1). Generally, ITNs in the lower group are made of tulle/PE fabrics, while the upper group is a mix of tulle/PE and traverse/PET. Consequently, these observations highlight that products cannot be judged on knitting pattern and polymer composition alone, and both polymers are capable of higher performance ITN products depending on quantitative design characteristics, including but not limited to, constituent yarn dimensions and mechanical properties, as well as the knitted fabric structure.

**Figure 1.**
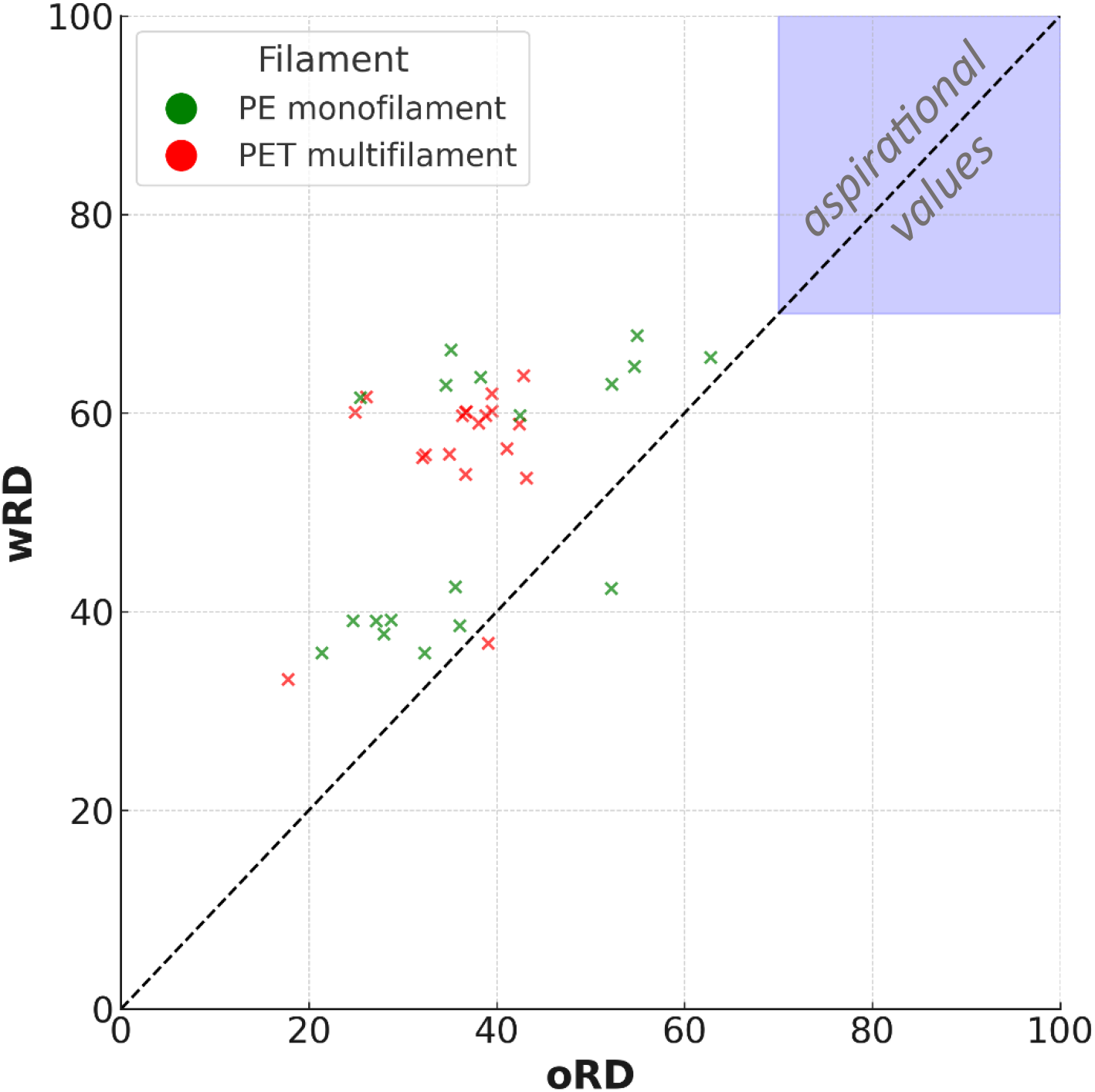
Comparison of oRD and wRD scores for different insecticide-treated nets (ITNs), categorized by tulle knitting pattern with polyethylene (PE) nets, and traverse knitting pattern with polyethyelene terephthalate (PET) nets

### Estimating median survival using oRD and wRD scores

When predicting ITN survival time, accurate predictions in the early stages of net lifespan (i.e. soon after distribution) is not as valuable as making accurate predictions lateron. In bivariable linear regression models, the power of the oRD to predict median survival at a given timepoint was highest for earlier study timepoints, when most nets are still present, with predictive power decreasing as the trial progresses (the R-square value decreased from 0.12 at 12-months to 0.00 at 36 months). In the same model, the wRD provided steady predictive power through the course of a trial, with the adjusted R-square value staying in the range 0.07 to 0.08 across the three survey rounds. The greater predictive power of the wRD compared to the oRD, particularly at later survey timepoints, came primarily from increasing the weighting of hole enlargement which is a strong differentiator between products. Differences in abrasion resistance between products was a negligible contributor to predictive power.

In the site-level analysis, a 10-point increase in wRD score was associated with a 3.9-month gain in median survival from first hanging (Figure 2). Furthermore, increasing wRD from 30 to 70, the approximate range for current products on the market, increases median survival by 13.7 months (95% CI: 3.1–24.2, *P* = 0.016, Figure 2). The wide error bars reflect country-level variation.

**Figure 2:**
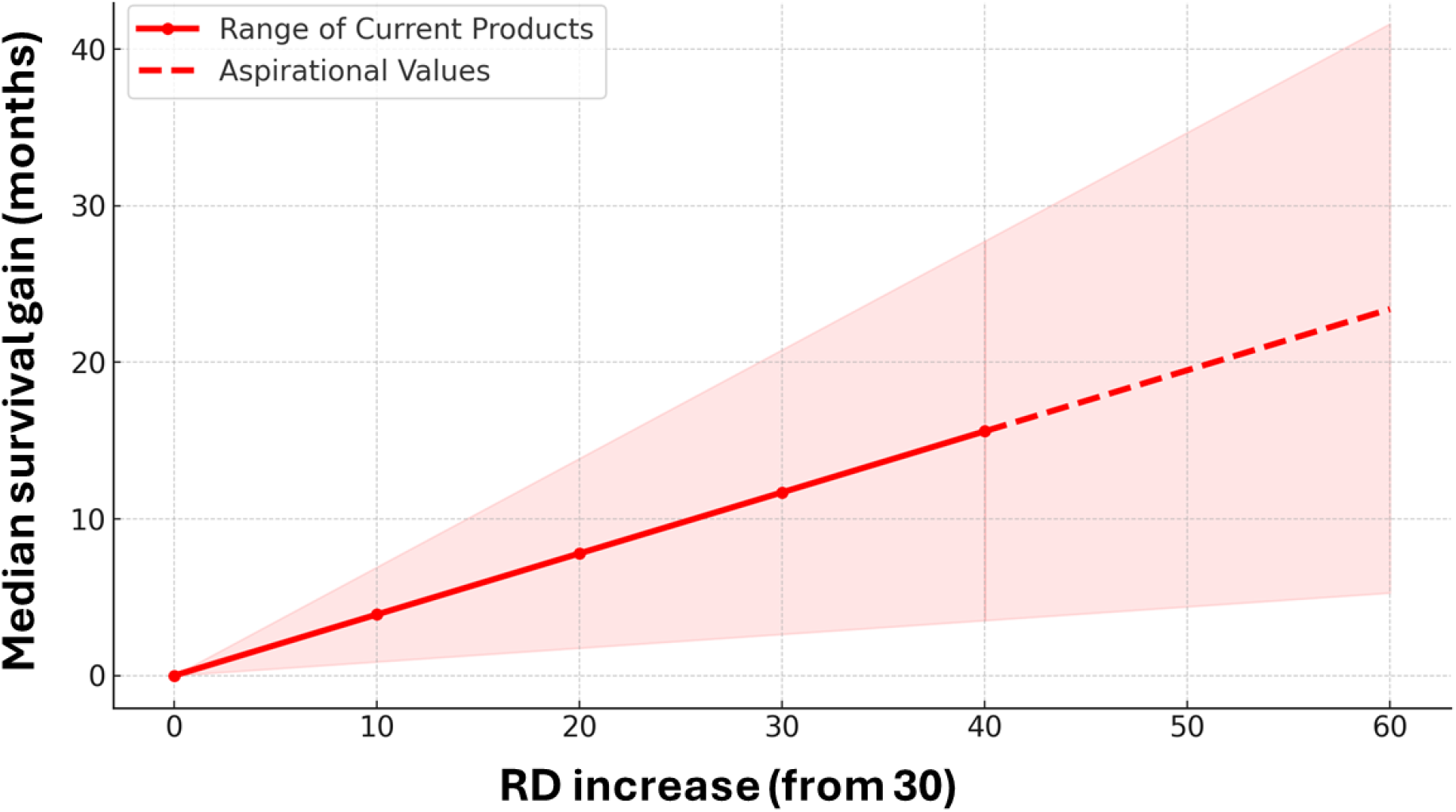
Visualization showing relative impact of increasing wRD score on median survival when increasing above a value of 30 (the lowest wRD score for ITN brands in our dataset) up to 70 (the highest wRD score for brands in our dataset). The dashed line indicates projected increases in survival by aspirational nets with physical durability beyond currently available ITN products. Shaded area indicates 95% confidence interval

Incorporating the Risk Index (RI) into the wRD model improved model fit, emphasizing the significant role of net handling, net care and repair and environmental factors in explaining variations in bednet lifespan across study areas (manuscript in preparation). Further investigation on the RI is underway to understand these contributing risk factors and enhance its usability, similar to the RD score.

ITN-level predicted survival from the Cox proportional hazards mixed effects regression model (adjusted for net use environment, net handling and net care attitude fixed effects, and country as a random effect) was plotted for wRD scores ranging from 38 to 68 (Figure 3). Within six months of ITNs being first hung, better survival is evident for products with higher wRD scores. The 50% survival point is reached approximately 1.2 years later for nets with a wRD score of 68 compared to those with a wRD score of 38, extending survival of hanging ITNs closer to the three-year target.

**Figure 3.**
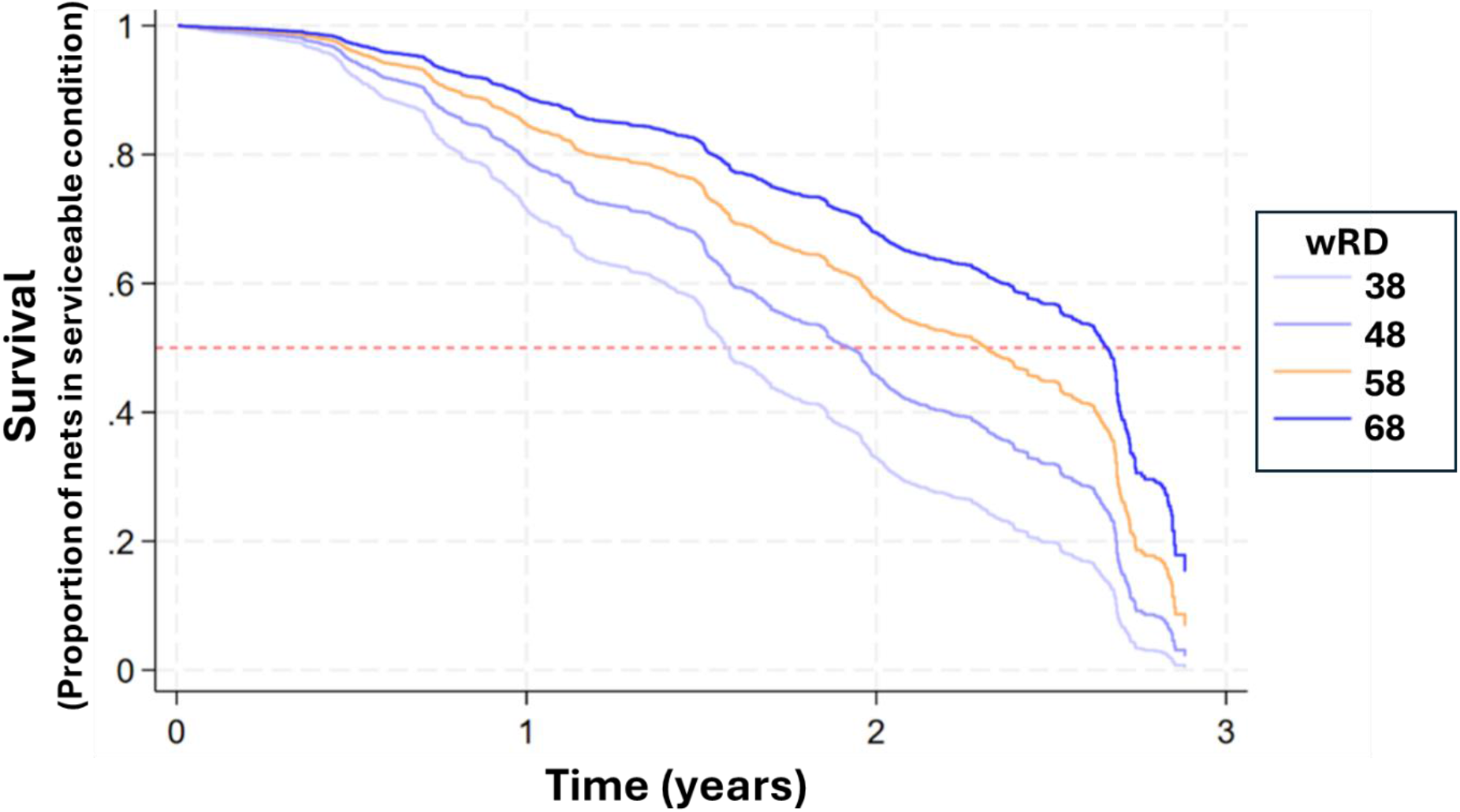
Predicted survival of ITNs with wRD values of 38, 38, 58, and 68, from ITN-level regression model. Dotted horizontal red line indicates 50% survival, at which point half of ITNs distributed are in serviceable condition

## Discussion

Here we show the value of a resistance to damage (RD) metric that equally values preventing holes from forming and minimising their propensity to enlarge, in predicting ITN lifespan under typical household use environments in sub-Saharan Africa. The wRD score is more predictive of net survivorship than the oRD score, due to increasing the weight of the resistance to hole enlargement variable. A 10-point increase in wRD score was associated with an estimated 3.9-month gain in median survival. This means, the expected difference in ITN lifespan between the lowest performing and highest performing nets on the market in 2020 (minimum: wRD 35– maximum: wRD70) is more than a year (13.75 months). This highlights the importance of both preventing holes from forming initially (e.g. due to snagging of the net fabric) and preventing these holes from enlarging, which is a mechanically distinct and equally important trait. We highlight that snag and bursting strength scores are very similar across the ITN product range, yet resistance to hole enlargement is a major differentiator between products. This differentiation is evident in the observation that wRD scores cluster into two groups, reflecting brands with higher and lower hole enlargement scores. However, we also draw attention to the observation that snag strength is poor across the entire product range, thus any future design that provided even moderate performance on this metric would compare well with current products on wRD. These findings support previous work by Kilian et al (2021) demonstrating that the components of the RD metric are predictive of operational lifespan [15].

We demonstrate that resistance to hole enlargement is a previously undervalued component in the oRD. By weighting hole formation (snag resistance and bursting strength) and hole enlargement equally in the rwRD score, we highlight that both traits should be a focus in the procurement and development of durable ITN designs. That resistance to hole enlargement is an important predictor of ITN lifespan is consistent with the observations by Wheldrake et al. [11] that most damage (by surface area) is caused by tearing of initially small holes. Furthermore, the finding that abrasion resistance alone is a negligible predictor of ITN lifespan is consistent with observations by Wheldrake et al. [11] in that <5% of all damage by surface area is caused by abrasion. Consequently, we summarise that the wRD is better aligned than the oRD to the damage mechanisms that ITNs face in the field.

The performance of engineered structures such as textile fabrics, depend on multiple factors, including their polymer composition, yarn and fabric structure, as well as dimensional and mechanical properties. It is noteworthy that all but one traverse/multifilament polyester ITN products exhibited excellent, near aspirational, resistance to hole enlargement. However, we emphasise that there are also examples of tulle/monofilament polyethylene nets with very good performance on the same metric. Consequently, the scoring of traverse/multifilament polyester products generally improved in the wRD model as compared to oRD. However, to attribute differences in the performance of a net solely to polymer content, e.g. polyester vs. polyethylene is misleading, as it ignores other important factors such as the yarn and fabric structure.

The ability of holes to enlarge following an initial yarn breakage is dependent primarily on the geometric structure (knitting pattern) and dimensional stability of the fabric, rather than its polymer composition. Unravelling and enlargement of an initial hole does not require additional yarn breakage. Changing the knitting pattern, and thereby, the frictional interaction of yarns within the knitted fabric structure to improve structural stability, is among the strategies that could be pursued to improve performance. Better hole enlargement resistance scores were also shown here to be positively correlated with mesh count, potentially due to the higher number of inter-yarn frictional contact points that are available to resist unravelling as the area between fibers gets smaller.

Investigating the physical design characteristics of ITNs in detail has highlighted two routes for innovation, i.e. snag resistance and hole enlargement resistance. Even the highest performing ITN products had poor snag strength relative to the aspirational target (Supplementary file 3). Clearly, major innovation in ITN design is needed. Knitted fabrics currently used to make ITNs are inherently susceptible to snagging on solid protuberances, leading to mechanical damage and yarn breakage. Consequently, snagging is the most common form of initial damage in nets retrieved from the field [11]. While holes associated with snagging are typically small, they are susceptible to enlarging over time, especially in net products with poor hole enlargement resistance. Further investigation to understand the mechanics of snagging and improve snag resistance is important. Hole enlargement resistance represents a completely different innovation path from snag strength, as it does not necessarily depend on filament breakage and requires distinct design considerations. To improve hole enlargement resistance, careful consideration should be given to the selection of knitting pattern and the tightness and dimensional stability of the fabric construction. These are all factors that can be modulated during textile manufacturing, but of course, potential trade-offs in terms of lower production speed, additional processing steps, higher fabric weight and therefore additional cost also need to be considered.

The long-term survivorship of ITN products depends on two basic factors. The first is the product design of nets in terms of their inherent physical durability, quantified by RD values. The second is the environment and the way in which the ITN is used, including net handling, net care and repair and environmental factors. While high-RD ITN products will withstand expected sources of mechanical damage in the household better than those with a lower value, certain household and environmental variables, as well as user care, expose the net to a heightened risk of damage. Consequently, even high-RD products would be expected to have short field life in very challenging conditions. However, as shown in our ITN-level regression, after adjusting for these known risks, ITN brands with wRD scores at the top of the current range were predicted to survive in serviceable condition for more than 14 months longer than brands with a wRD score and the bottom of the current range, on average. Understanding how much additional survival gain could be achieved from ITNs with even higher wRD scores is an important unanswered research question.

The ITN products investigated here fall well below the aspirational value of wRD score of 100[10, 12], particularly on snag strength, indicating room for considerable improvement in ITN design in terms of physical durability. Consequently, there is a need to explore fabric design characteristics that could result in a higher wRD score. For manufacturers, this requires new incentives to innovate, as at present there is no clear market willing to pay the costs associated with the development and manufacturing of more durable nets. Funding for malaria control is limited [1] and a case for value-for-money, rather than cost, needs to be made. The potential cost effectiveness afforded by longer-lasting nets may be considered by ITN procurers and country programmes. In addition, sufficient incentive must be afforded to manufacturers to be able to explore improved and innovative durability design. These innovations must go together with a renewed focus on net care to give these products the best opportunity to perform as expected in the field.

## Conclusion

ITN products with higher wRD scores are projected to provide longer field life, with an increase from 35 to 70 estimated to provide 13.65 additional months in serviceable condition. This can have immediate implications on ITN procurement decisions to encourage the purchase of nets that will last longer in the field and it provides direction on innovation to improve net retention. This study underlines the importance of determining the wRD score of an ITN product during ITN prequalification, to ensure nets can provide long-term physical protection in the field. The wRD score demonstrates greater accuracy in predicting net survivorship in the field compared to the oRD score. Therefore, we propose adopting the wRD score as the new RD score standard and encourage its endorsement, standardisation and uptake in procurement decisions by donors and country programmes. By giving equal weight to resistance against hole formation and hole enlargement, the wRD score highlights that preventing small holes from expanding is just as crucial as preventing their formation. To address this, alternative knitting patterns and new ways of producing nets could enhance resistance to hole enlargement, while innovations targeting snag resistance can further boost net physical durability. There is a pressing need for genuine innovation to produce more physically durable nets, and recognizing the long-term value of physical durability can justify the inevitable higher initial costs.

## Declarations

### Authors’ contribution

FM, SP, SJR and AS designed the study protocol. FM, SP, JAT, MW and ES, AW and FJ participated on data analysis. FM, SP, JAT drafted the manuscript. MW, ES, AK, HK, CF, SJR and AS critically reviewed the manuscript. All authors read and approved the final manuscript.

## Acknowledgments

The authors would like to acknowledge

## Availability of data and Materials

Data supporting the conclusions of this article are included within the article. Datasets are available on reasonable request to NIRI, UK.

## Ethical approval

Not applicable

## Consent for publication

Not applicable

## Supplementary files

**Supplementary file 1:**
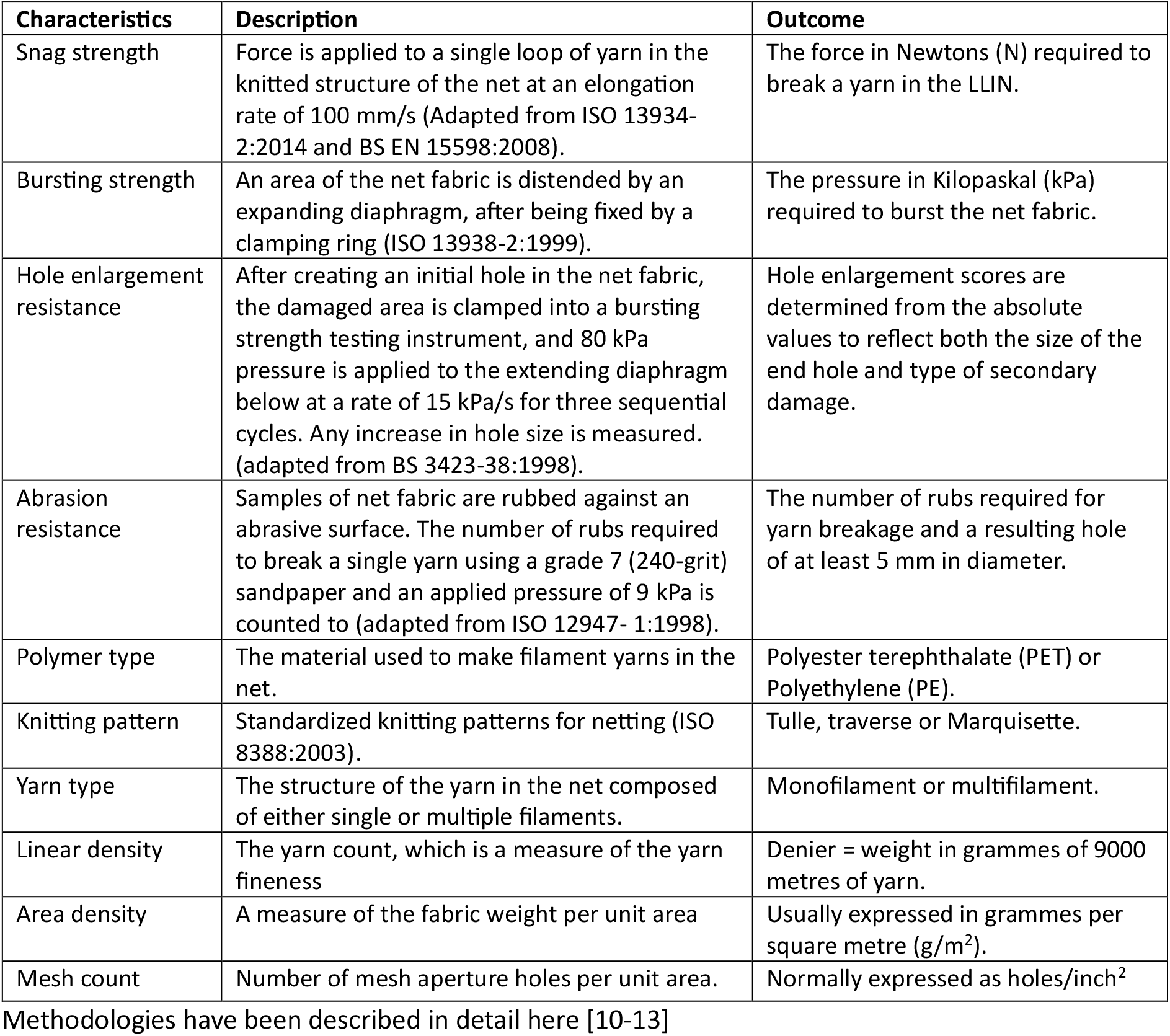
List of physical characteristics of interest.

**Supplementary file 2:**
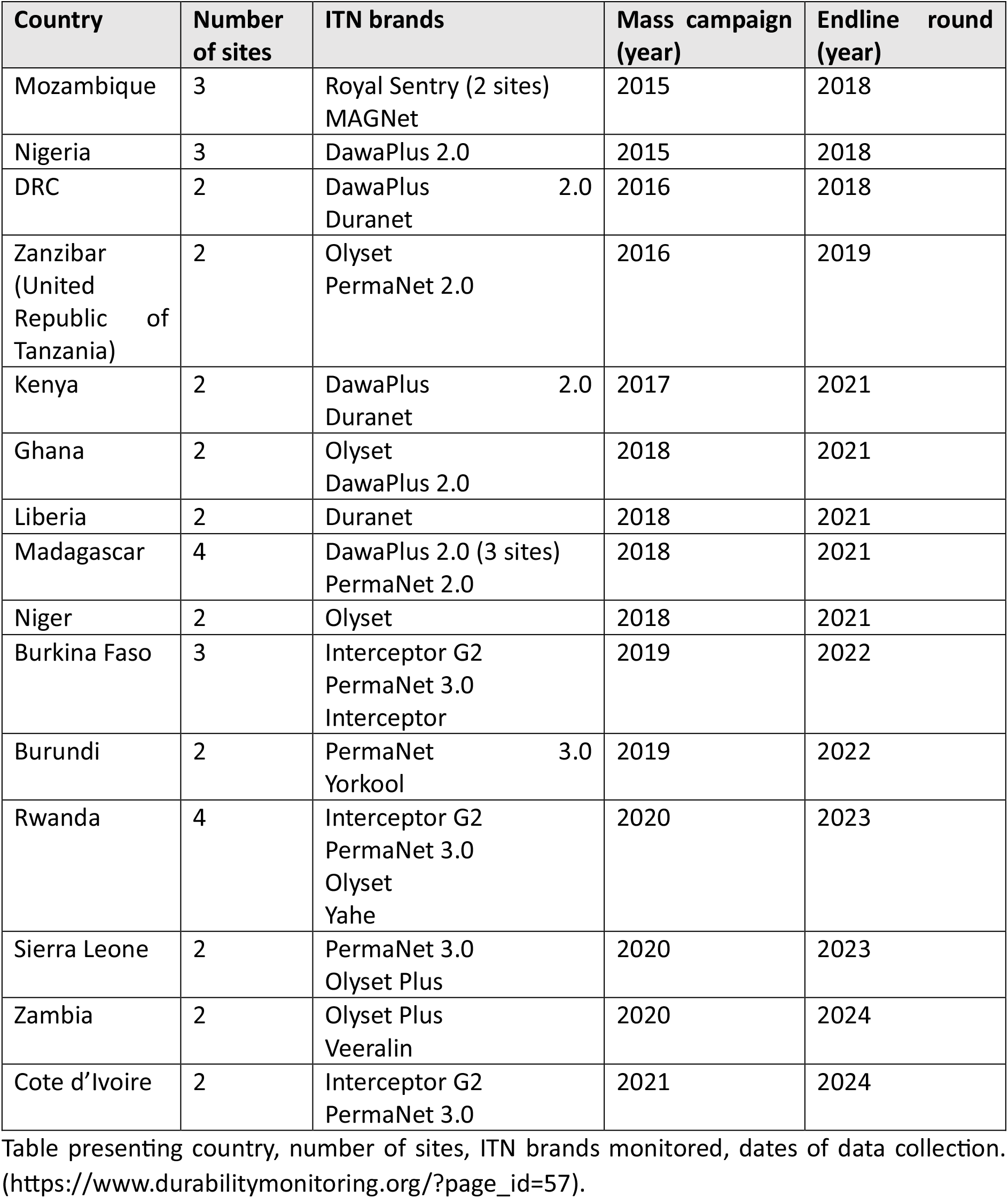
Summary of durability monitoring data collection

**Supplementary file 3:**
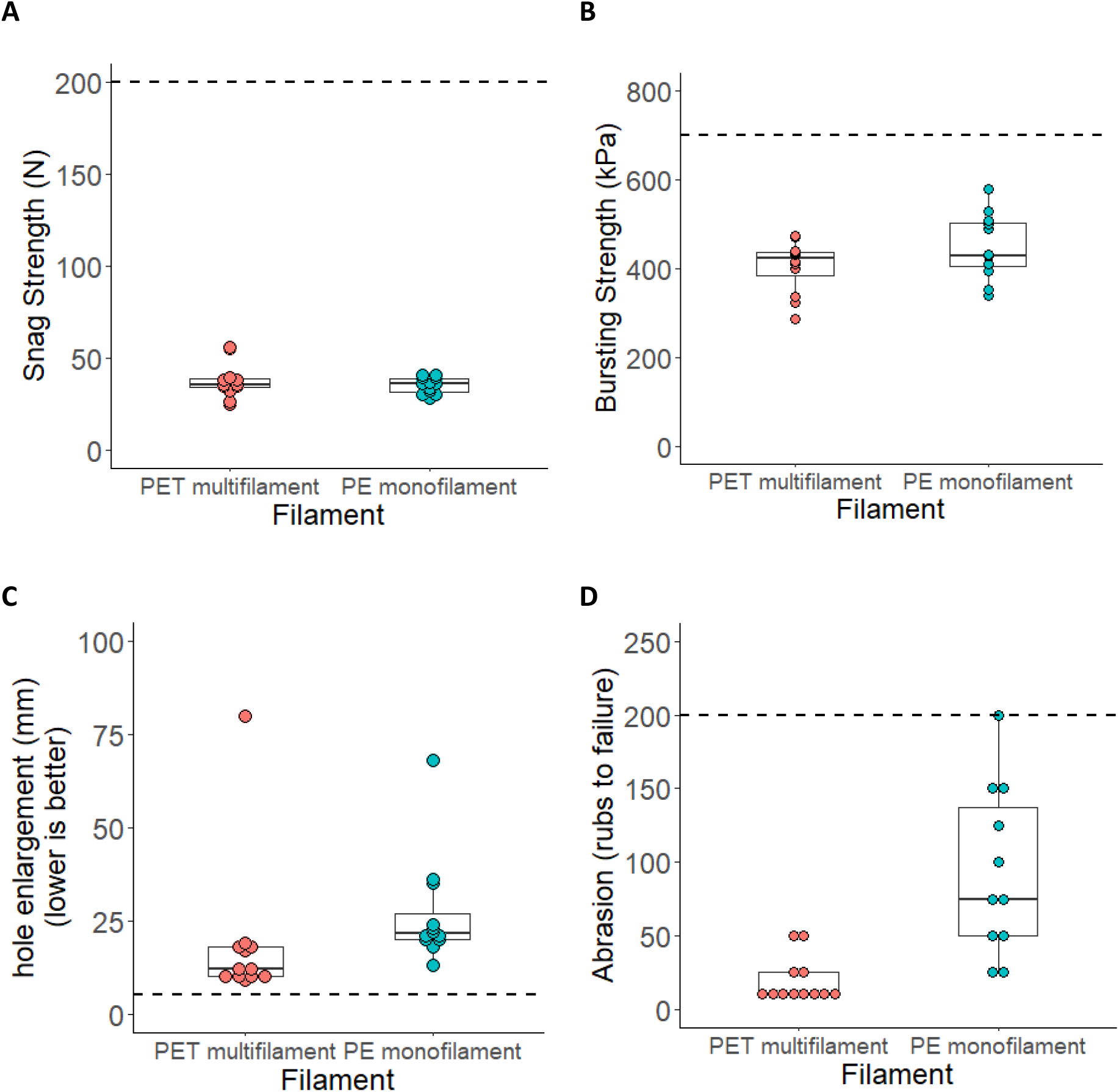
Physical characteristics of nets in relation to their polymer composition and filament types, snag strenghth, bursting strength, hole enlargement resistance and abrasion resistance. Dotted, horizontal line indicates aspirational score.

**Supplementary file 4:**
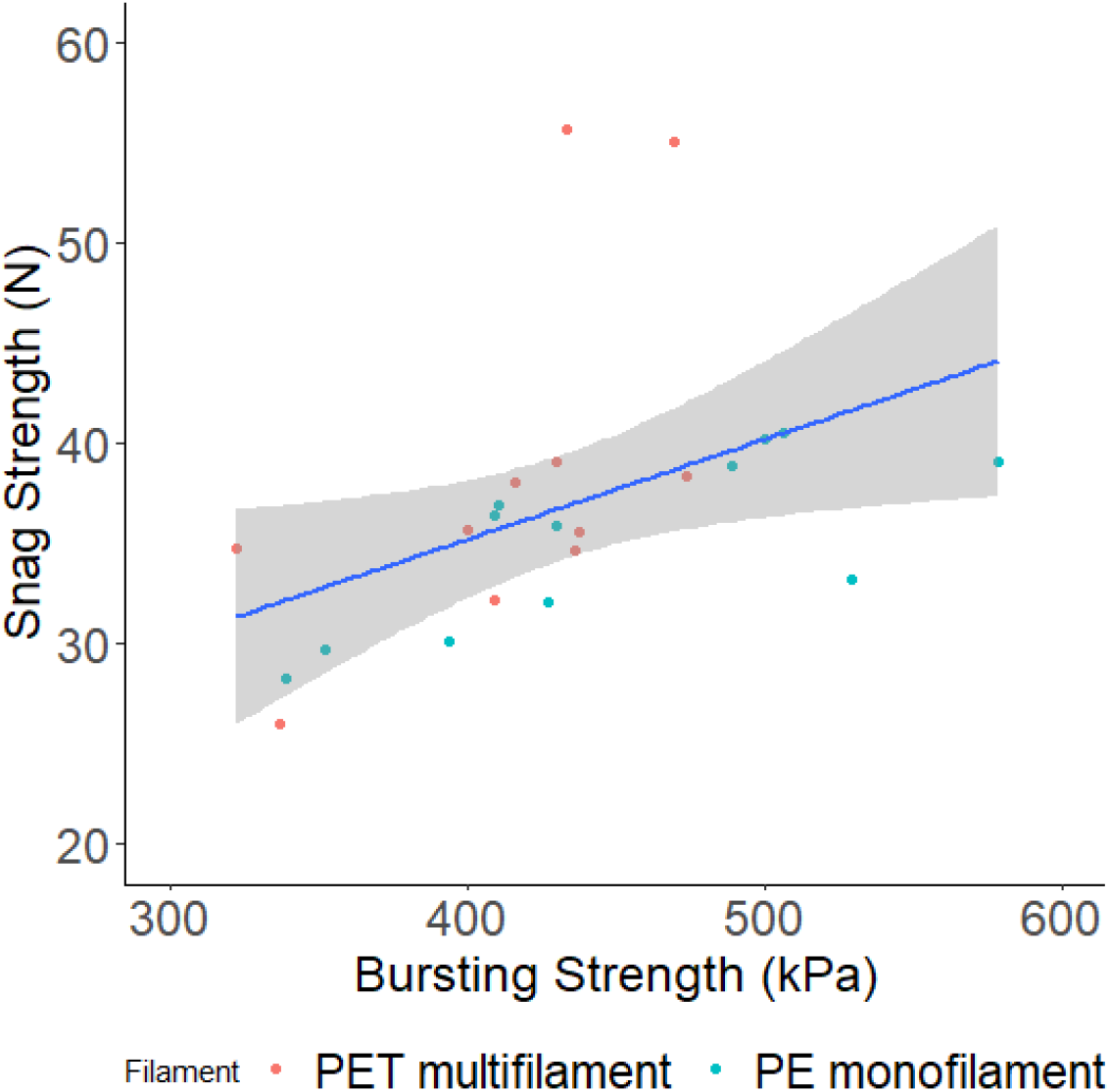
Correlation between Snag strength and Bursting strength. N= Newtons, kPa = kilopascals. The snag strength and bursting strength are well correlated with each other, with a Spearman correlation of 0.661.

**Supplementary file 5:**
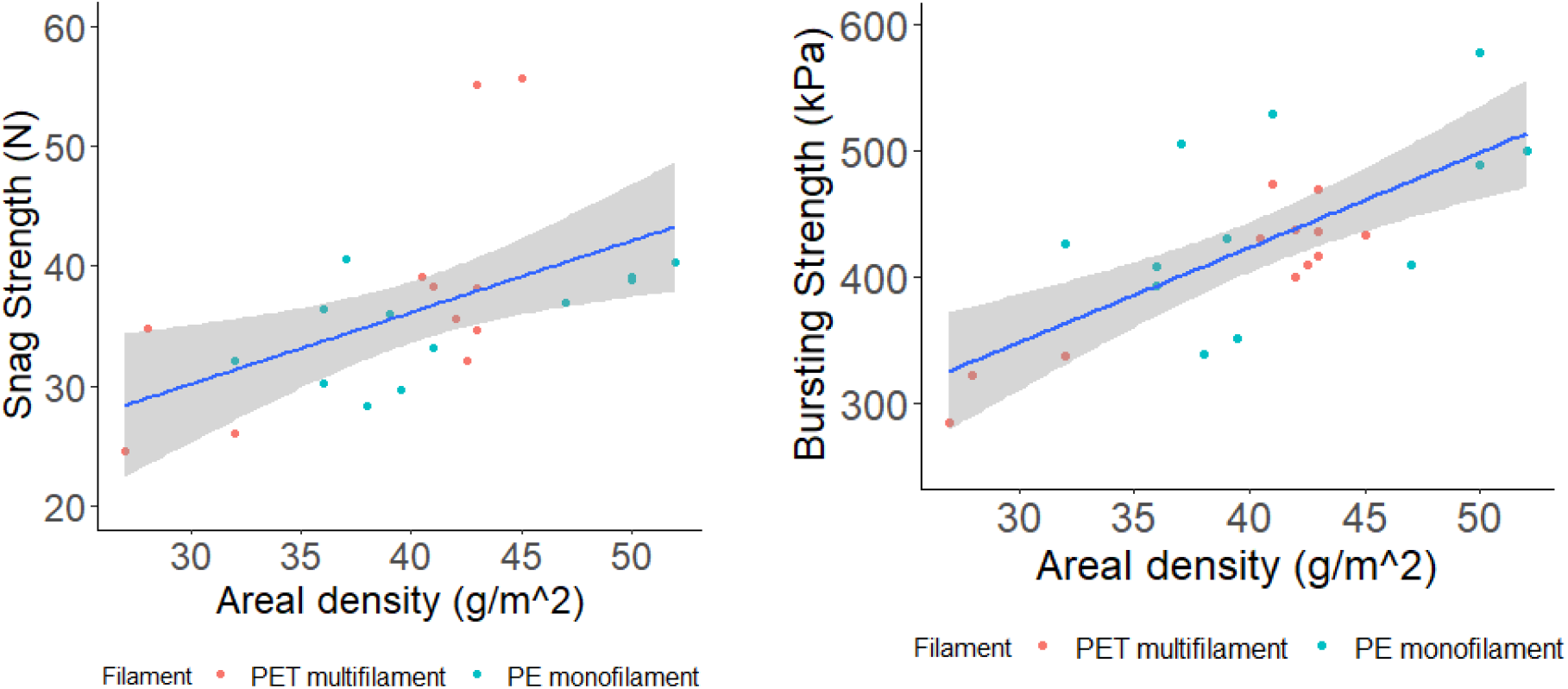
Relationship between area density and snag/burst strength

**Supplementary file 6:**
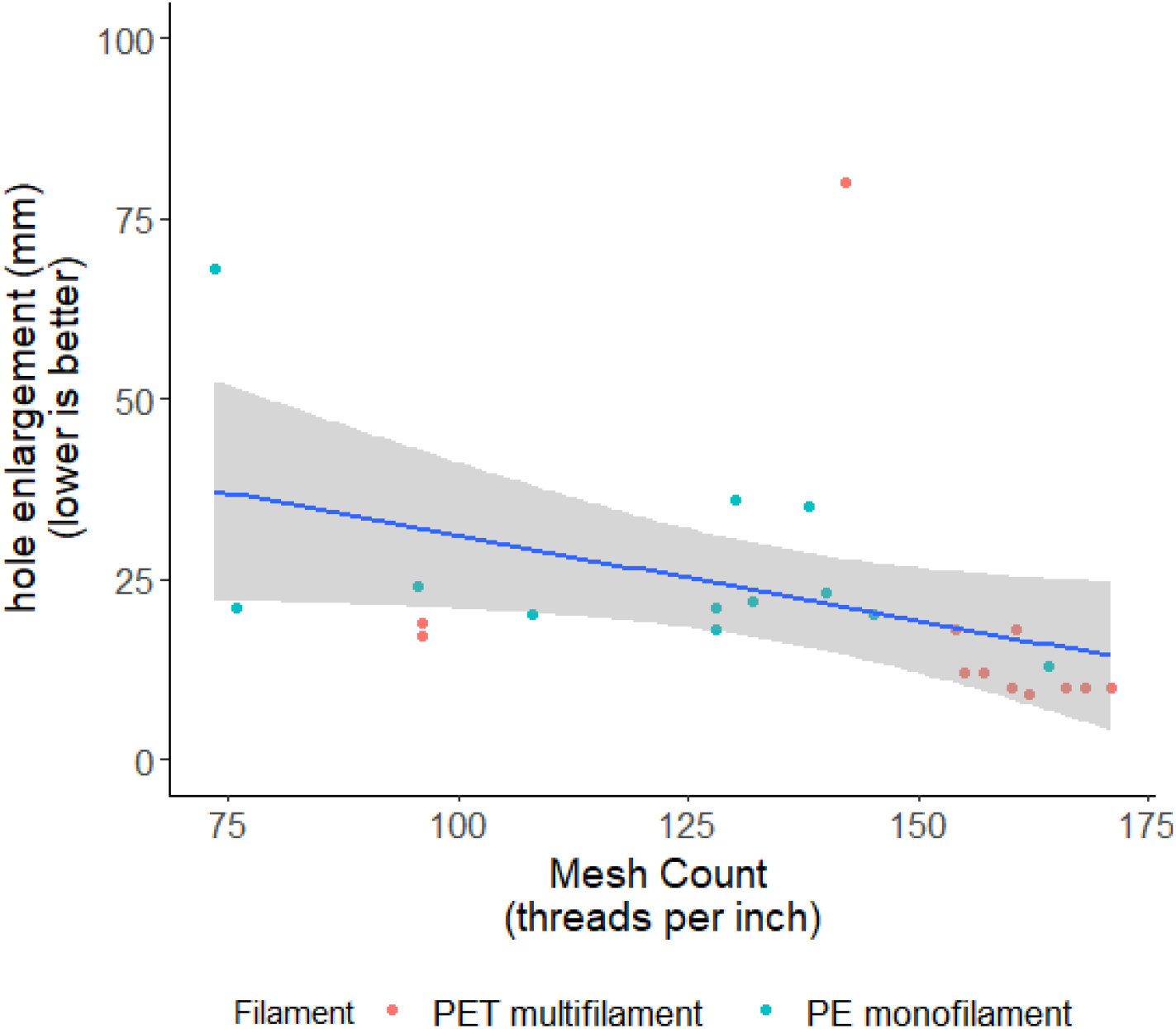
Relationship between mesh count and hole enlargement

